# FANCJ promotes PARP1 activity during DNA replication that is essential in BRCA1 deficient cells

**DOI:** 10.1101/2024.01.04.574095

**Authors:** Ke Cong, Nathan MacGilvary, Silviana Lee, Shannon G. MacLeod, Jennifer Calvo, Min Peng, Arne Nedergaard Kousholt, Tovah Day, Sharon B. Cantor

## Abstract

Single-stranded DNA gaps are postulated to be fundamental to the mechanism of anti-cancer drugs. Gaining insights into their induction could therefore be pivotal for advancing therapeutic strategies. For poly (ADP-ribose) polymerase inhibitors (PARPi) to be effective, the presence of FANCJ helicase is required. However, the relationship between FANCJ dependent gaps and PARP1 catalytic inhibition or trapping—both linked to PARPi toxicity in BRCA deficient cells—is yet to be elucidated. Here, we find that the efficacy of PARPi is contingent on S-phase PARP1 activity, which is compromised in FANCJ deficient cells because PARP1, along with MSH2, is “sequestered” by G-quadruplexes. PARP1’s replication activity is also diminished in cells missing a FANCJ-MLH1 interaction, but in such cells, depleting MSH2 can release sequestered PARP1, restoring PARPi-induced gaps and sensitivity. Our observations indicate that sequestered and trapped PARP1 are different chromatin-bound forms, with FANCJ loss increasing PARPi resistance in cells susceptible to canonical PARP1 trapping. However, in BRCA1 null cells, the loss of FANCJ mirrors the effects of PARP1 loss or inhibition, with the common detrimental factor being the loss of PARP1 activity during DNA replication, not trapping. These insights underline the crucial role of PARP1 activity during DNA replication in BRCA deficient cells and emphasize the importance of understanding drug mechanisms for enhancing precision medicine.

## INTRODUCTION

Poly(ADP-ribose) polymerase inhibitors (PARPi) are effective in treating cancers with mutations in the hereditary breast and ovarian cancer genes, BRCA1 or BRCA2 (BRCA)^1,2^. The underlying PARPi sensitization mechanism remains unclear given that PARP1 has a range of distinct functions beyond homologous recombination (HR). The multifaceted roles of PARP1 span from excision repair, including single-strand break repair, base excision repair and nucleotide excision repair, to mediating both classical and alternative non-homologous end joining, to the regulation of replication fork dynamics and DNA end resection activities^3–9^. Moreover, PARPi toxicity is thought to stem from chromatin associated “trapped” PARP1. The inhibitors disrupt PARP1’s catalytic activity, which in turn reduces the auto-poly(ADP-ribosyl)ation (auto-PARylation) that releases PARP1 from chromatin^10,11^. Thus, the collision of replication forks with either unrepaired single-strand breaks or trapped PARP1 complexes has been proposed to induce DNA double-strand breaks that necessitate BRCA1 function in HR^1,2^. Additionally, PARP1 functions in DNA replication as a backup to canonical lagging strand synthesis^12^. Upon detection of un-ligated lagging strands, PARP1 creates poly(ADP-ribose) (PAR) covalent attachments on itself and XRCC1 and LIG3 to promote gap filling. Accordingly, disruption of lagging strand synthesis and the trapping of PARP1 at unligated lagging strands could underlie toxicity^13^.

We have reported that PARPi-induced single-stranded DNA (ssDNA) gaps are closely aligned with PARPi toxicity. Gaps as a determinant of PARPi response were in part highlighted by the comparison between cells deficient in the hereditary breast and ovarian cancer genes BRCA1 or FANCJ. Cells deficient in either gene display similar defects in HR and fork protection^14–16^. By contrast, BRCA1 and FANCJ deficient cells differ in their relationship to PARP1. PARPi induced few replication gaps and little sensitivity in FANCJ deficient cells as compared to high gap accumulation and synthetic lethality with BRCA deficiency. Moreover, PARP1 activity during S phase is abnormally low in FANCJ deficient cells but elevated in BRCA1 deficient cells. Finally, while replication speed is also distinct between FANCJ versus BRCA1 deficient cells, a series of cell models revealed that speed can be uncoupled from PARPi toxicity^17^. The contrast between BRCA1 and FANCJ is further supported by a genetic interaction network of PARPi response^18^. Notably, FANCJ loss as compared to other BRCA-Fanconi anemia genes confers the weakest PARPi sensitization phenotype, indicating again the unique role of FANCJ. Together, these findings suggest that PARP1 replication activity is central to the induction of PARPi induced gaps and sensitivity, but how FANCJ confers these outcomes is unclear.

We hypothesized that FANCJ loss could impact PARP1 activation via changes in the replisome composition and/or DNA secondary structures. G-quadruplexes (G4s) form in lagging strands and are elevated in the replisome of FANCJ deficient cells^19–21^. G4 processing by FANCJ requires its helicase and translocase activities as well as the integrity of lysines 141 and 142 that bind G4s^22,23^. These lysine residues also mediate an interaction between FANCJ and the mismatch repair (MMR) protein MLH1 that is required for replication stress recovery. Restart in cells lacking the FANCJ-MLH1 interaction can be restored by depletion of the MMR protein MSH2, suggesting that dysregulated MMR limits replication in FANCJ deficient cells^24,25^. Intriguingly, beyond its role in correcting post-replication DNA mismatches, MMR has heightened activity on lagging strands^19–21^. Restricting MMR during replication could potentially be the mechanism by which FANCJ activates PARP1’s backup function on the lagging strand. This may clarify why, in FANCJ deficient cells, there is a modest presence of PARP1 activity, PARPi-induced ssDNA gaps and sensitivity^17^.

Here, we uncover that PARP1 function during DNA replication is disrupted in FANCJ deficient cells even though PARP1 chromatin loading and activation by DNA damaging agents remain intact. Our data support a model in which FANCJ dismantles replisome-associated MSH2-bound G4s that limit ssDNA and PARP1 activation. This model is supported by the finding that MSH2 depletion re-establishes PARP1 activity along with PARPi-induced gaps and sensitivity in cells that lack the FANCJ-MLH1 interaction. Based on the discovery that BRCA1 deficient cells have elevated PARP1 activity and extreme sensitivity to any perturbation that disrupts PARP1 replication activity, we can further draw the conclusion that loss of PARP1 activity as opposed to canonical PARP1 trapping is the primary cause of cytotoxicity in these cells. Given that PARP1 trapping is toxic in other contexts, our findings underscore the importance of understanding drug action in precision medicine, to facilitate optimal drug combinations and prevent resistance that could evolve from low PARP1 S phase activity and PARP1 trapping.

## RESULTS

### FANCJ promotes PARP1 activity during DNA replication and limits MSH2 loading at the replisome

We previously found minimal PARPi sensitivity in FANCJ deficient cells including human retinal pigment epithelial 1 (RPE1), osteosarcoma (U2OS) and immortalized human embryonic kidney 293T cells^17^. Further characterization of FANCJ knockout (KO) RPE1 cells revealed aberrantly low PARP1 activity as measured by PARylation (PAR) in contrast to BRCA1 KO cells in which PAR was higher than wild-type (WT) controls^17^. Thus, we sought to determine if PAR was also low in the FANCJ KO U2OS and 293T cell systems. To identify the endogenous PARP activity without external genotoxic stress as previously reported^12^, we utilized an inhibitor of PARG, an enzyme primarily accountable for poly(ADP-ribose) removal, and observed elevated poly(ADP-ribose) or PARylation that correlated with replication^17^. Similar to the RPE1 cells, we observed low PAR in the FANCJ null U2OS cells as measured by immunoblot following PARGi incubation (**Fig. 1a-1b**). Low PAR was also detected in FANCJ null 293T cells by immunoblot (**Fig. 1c-1d**). Similar to our prior findings in RPE1 cells, the FANCJ KO U2OS or 293T cells activated PARP1 following treatment with DNA damaging agents, methylmethane sulfonate (MMS) and hydrogen peroxide (H_2_O_2_) (**Supplementary Fig. 1a, b**)^17^ demonstrating there was not an intrinsic defect in PARP1 enzyme activity and that FANCJ was not required for PARP1 activation in response to DNA damage. As found in RPE1 cells, we also observed that the lower PARP1 activity was unique to S phase or actively replicating U2OS cells positively incorporating the nucleotide analogue 5-ethynyl-2′-deoxyuridine (EdU) as measured by immunofluorescence (**Fig. 1e**).

**Fig. 1:**
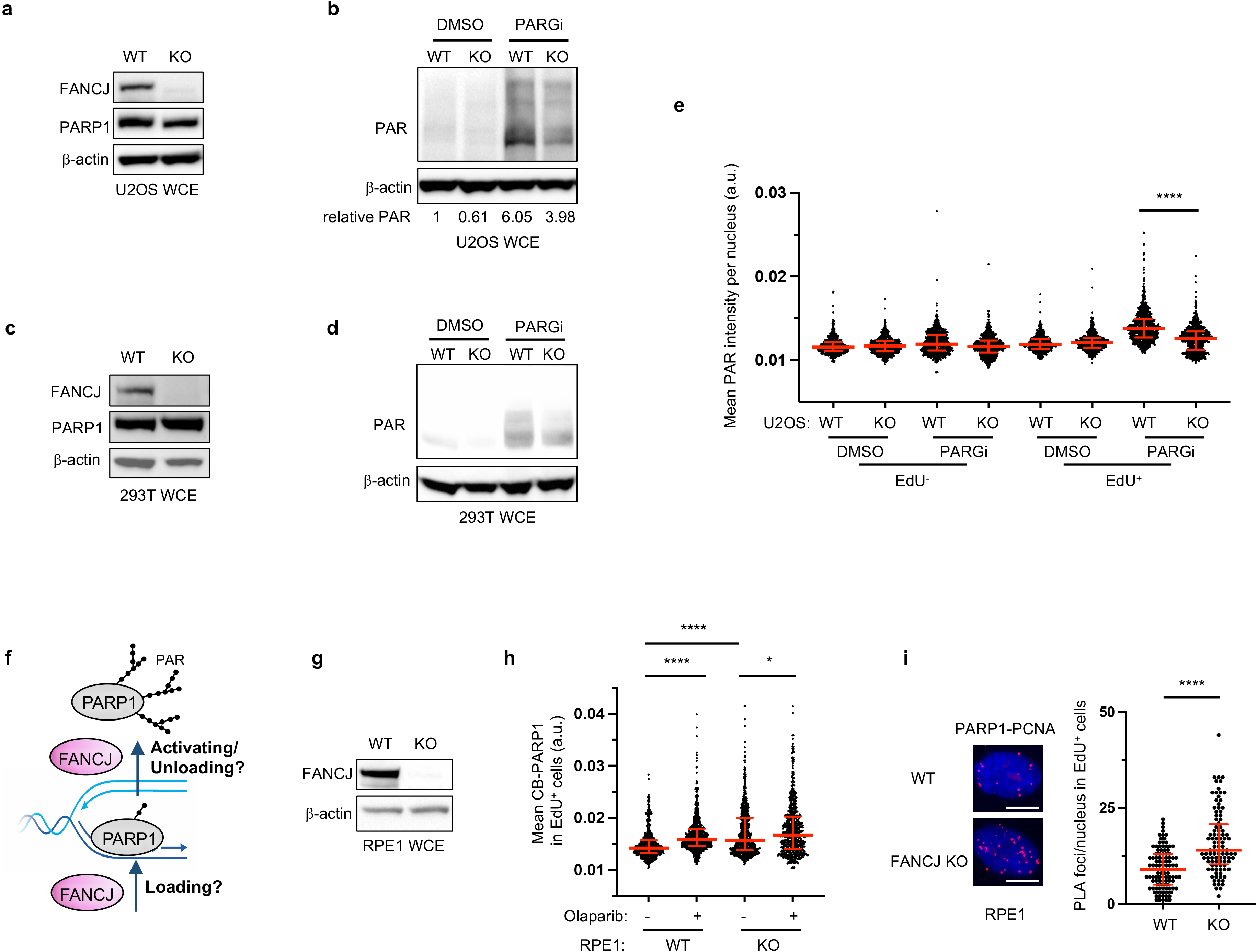
Modest PARPi sensitivity of FANCJ deficient cells is associated with low S phase PAR, enhanced MSH2 and PARP1 chromatin loading. **a** Representative western blot (WB) for the whole cell lysates (WCE) showing indicated proteins in untreated WT vs FANCJ KO U2OS cells. **b** Representative WB for the whole cell lysates showing the PAR formation in indicated U2OS cells treated with DMSO or PARG inhibitor (PARGi, 10 µM) for 40 min prior to harvesting to block PAR removal. Quantification of mean relative PAR from three biological replicates normalized to β-actin shown. **c** Representative WB for the whole cell lysates showing indicated proteins in untreated WT vs FANCJ KO 293T cells. **d** Representative WB showing the PAR formation in indicated 293T cells treated with DMSO or PARG inhibitor (PARGi, 10 µM) for 40 min prior to harvesting to block PAR removal. **e** Quantification of mean PAR intensity per nucleus for WT and FANCJ KO U2OS cells treated with DMSO or PARGi (10 µM, 30min) together with EdU incubation. **f** Schematic showing how FANCJ impacts PAR formation during replication: could FANCJ impact PARP1 loading and/or releasing during replication? **g** Representative WB for the whole cell lysates showing indicated proteins in untreated WT vs FANCJ KO RPE1 cells. **h** Quantification of chromatin-bound PARP1 (CB-PARP1) for RPE1 WT and FANCJ KO with or without Olaparib treatment (10 µM, 2hrs), with EdU incubated at the final 30min. For **e** and **h**, EdU-positive or EdU^+^ cells were gated according to positive EdU incorporation. Each dot represents one cell. At least 300 cells are quantified from 3 biological independent experiments (n = 3). Red bars represent the median ± interquartile range. All statistical analysis according to Kruskal-Wallis test, followed by Dunn’s test. **i** PARP1-PCNA proximity ligation assay (PLA) in untreated RPE1 WT vs FANCJ KO cells, with 10 µM EdU incubated for 20 mins. Dot plot shows the number of foci and the median ± interquartile range for at least 100 cells from 3 biological independent experiments (n=3). Scale bars, 10 µm. Statistical analysis according to Mann-Whitney test.

The low S phase PARP1 activity in cells without the DNA helicase/translocase activity of FANCJ could result from either a failure to efficiently load or unload PARP1 from chromatin (**Fig. 1f**). Thus, we sought to address the impact of FANCJ on replication associated PARP1. We first re-examined our prior iPOND (isolation of proteins on nascent DNA) data. As compared to WT, FANCJ KO 293T cells had enriched replisome and overall chromatin associated PARP1^16^. Consistent with this finding, we observed greater chromatin bound (CB) PARP1 in the FANCJ null RPE1 cells that was selective to EdU positive cells and was only modestly increased following PARPi treatment (**Fig. 1g, h**, **Supplementary Fig. 1c**). The PARP1 replisome enrichment in FANCJ KO RPE1 cells was also observed by proximity ligation assay (PLA) compared to a PCNA-PCNA PLA control (**Fig. 1i and Supplementary Fig. 1d**). Together, these findings suggest that FANCJ is dispensable for PARP1 chromatin localization but rather critical for its activation and/or unloading (**Fig. 1f**).

To understand how FANCJ could promote PARP1 unloading and/or S phase activation, we considered that in addition to PARP1, the MMR proteins MSH2 and MSH6, that form the MutSα complex, were elevated in the replisome of FANCJ KO 293T cells as suggested in our previous iPOND study^16^. We also found enriched MSH2-PCNA PLA signal and overall enriched CB-MSH2 in FANCJ KO RPE1 as compared to WT cells (**Fig. 2a and Supplementary Fig. 2a**). Furthermore, we observed a more pronounced MSH2-PARP1 PLA signal in FANCJ KO RPE1 cells (**Fig. 2b**) suggesting the proximity between PARP1 and MSH2 were aberrantly elevated and proximal to the replisome of FANCJ null cells. These findings in conjunction with our prior research linking FANCJ to MMR^24^ provided evidence that PARP1 and the MMR pathway are dysregulated in FANCJ deficient cells.

**Fig. 2:**
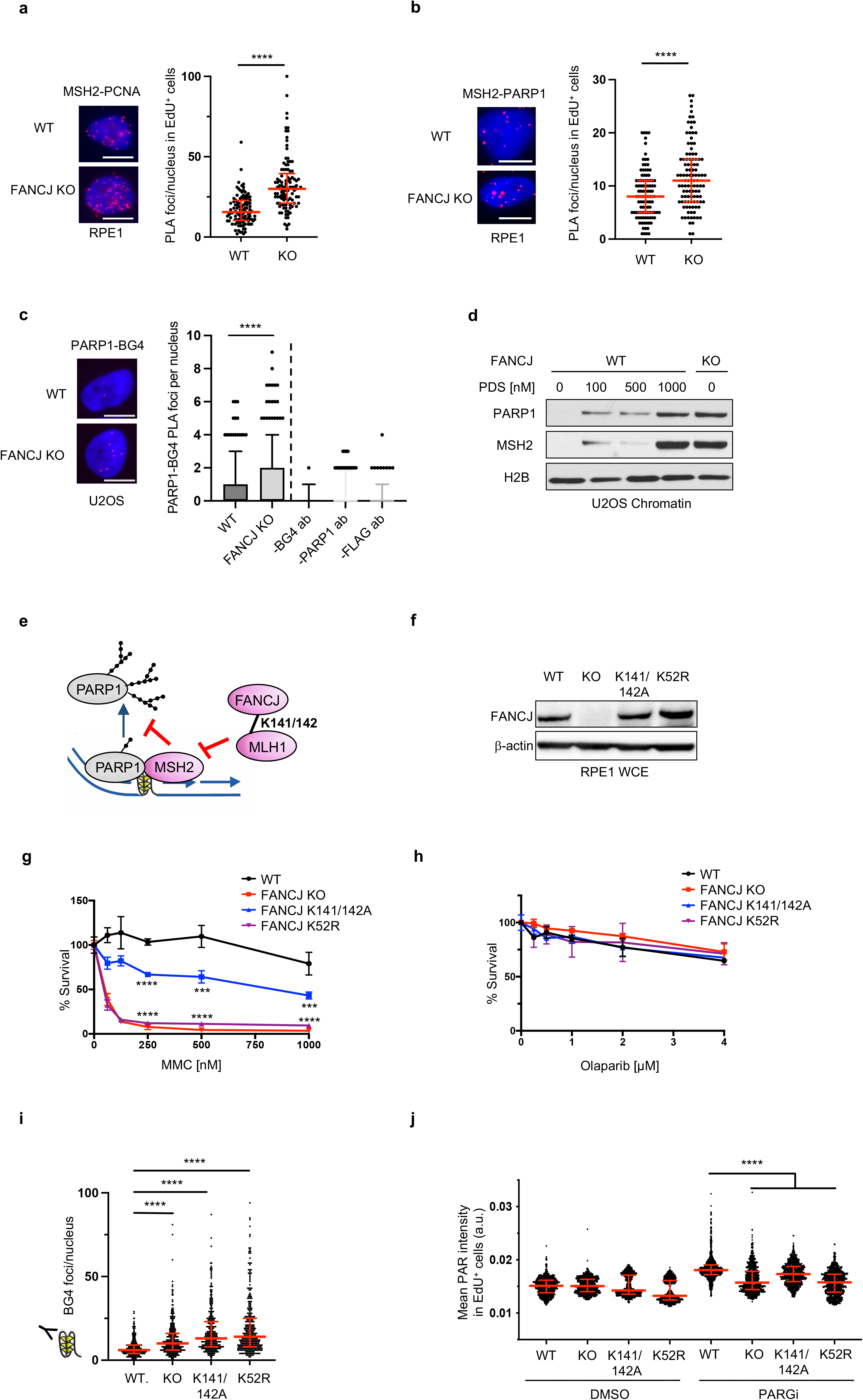
S phase PAR requires FANCJ helicase and MLH1 binding activities. **a** MSH2-PCNA PLA in untreated RPE1 WT vs FANCJ KO cells, with EdU incubated for 20 mins. **b** MSH2-PARP1 PLA in untreated RPE1 WT vs FANCJ KO cells, with EdU incubated for 20 mins. **c** PARP1-BG4 PLA in untreated U2OS WT vs FANCJ KO cells. **d** Representative WB of chromatin bound PARP1 and MSH2 in PDS treated U2OS WT cells compared to untreated WT and FANCJ KO cells. **e** Model showing MSH2 could limit PARP1 activation and be regulated by FANCJ-MLH1 interaction. **f** Representative WB analysis for FANCJ in the whole cell lysates of FANCJ knock-in mutant cells. **g** Cell survival assays for indicated cells under increasing concentrations of mitomycin C (MMC). Data represent the mean percentage ± SD of survival for each dot. **h** Cell survival assays for indicated cells under increasing concentrations of Olaparib. Data represent the mean percentage ± SD of survival for each dot. For **g** and **h**, significance was determined by one-way ANOVA followed by Dunnett’s test comparing FANCJ WT to mutant cells. **i** Quantification of G4 (BG4) foci/nucleus in RPE1 mutant cells under untreated growth conditions. **j** Quantification of PAR after 30min DMSO or PARGi (10 µM) treatment with EdU in EdU^+^ RPE1 WT, FANCJ KO, FANCJ K141/142A and K52R cells. For PLA assays above, dot plot shows the number of foci and the median ± interquartile range (**a** and **b**), or is represented as box-and-whisker with 5-95 percentiles (**c**) for at least 100 cells from 3 biological independent experiments (n=3). Scale bars, 10 µm. All statistical analysis according to Mann-Whitney test. For other quantification above, EdU^+^ cells were gated according to positive EdU incorporation. Each dot represents one cell. For **i**, At least 200 cells are quantified from n = 2. For **j**, at least 300 cells are quantified from n = 3. Red bars represent the median ± interquartile range. All statistical analysis according to Kruskal-Wallis test, followed by Dunn’s test.

Given that MMR and PARP1 bind G4 structures that form in the replisome of FANCJ deficient cells^26–33^, we sought to test if the proximity of PARP1 to G4s was also enhanced. First, we confirmed that the BG4 antibody detected greater signal in the RPE1 and U2OS FANCJ null cells as well as in WT cells following treatment with the G4 stabilizing agent, pyridostatin (PDS) (**Supplementary Fig. 2b, c**). Consistent with prior studies^26,27^, we find through proximity ligation analysis that PARP1 and G4 DNA are in close proximity (**Fig. 2c**). Moreover, we observe significantly increased PARP1-G4 PLA signal in FANCJ KO relative to WT cells (**Fig. 2c**). We also observe increased chromatin bound PARP1 and MSH2 in FANCJ null cells or in WT cells treated with PDS suggesting that G4s are bound by PARP1 and MMR proteins (**Fig 2d**, **Supplementary Fig. 2d**). Collectively, these findings demonstrate that FANCJ restricts the proximity of MSH2 with PARP1 and PARP1 with G4s. Thus, we hypothesized that the accumulation of PARP1/MSH2 bound G4s could limit PARP1 activation in FANCJ null cells (**Fig. 2e**).

### FANCJ helicase activity and interaction with MLH1 are required for PARP1 activity during DNA replication

FANCJ helicase activity resolves G4s and FANCJ lysines K141/142 bind G4s^23^ as well as mediating an interaction between FANCJ and the MMR protein MLH1^25^. Thus, we expected that mutations disrupting either function would elevate G4s and in turn reduce PARP1 activity. To test this prediction, we generated RPE1 cells with lysine 52 converted to arginine (K52R) to inactivate FANCJ ATPase and helicase activity^34^ and lysines 141 and 142 of FANCJ converted to alanine (K141/142A). We observed that the mutant proteins were expressed at similar levels to wild-type FANCJ (**Fig. 2f**). While the mutants enhanced sensitivity to the crosslinking agent, mitomycin C (MMC), PARPi resistance remained similar to the wildtype and null lines (**Fig. 2g, h**). In addition, we employed a re-expression strategy using either FANCJ KO or Fanconi anemia (FA) patient FANCJ deficient (FA-J) cells showing expected MMC sensitivity (**Supplementary Fig. 2e-h**) and PARPi resistance (**Supplementary Fig. 2i**). We also confirmed that the KO, knock-in (KI) RPE1 or complemented RPE1 or FA-J cells displayed a significant restart defect following release from aphidicolin (APH) (**Supplementary Fig. 2j,k**) or hydroxyurea (HU) (**Supplementary Fig. 2l,m**)^25^. Furthermore, as expected, we observed greater G4 accumulation by BG4 antibody immunofluorescence in the KO and mutant lines as compared to WT cells (**Fig. 2i**). Finally, examination of PARP1 activity in unperturbed S phase revealed that cell lines lacking FANCJ, its helicase activity, or MLH1 binding had low S phase PAR compared to FANCJ proficient cells (**Fig. 2j and Supplementary Fig. 2n**). Collectively, these findings suggest that G4s have the potential to restrict PARP1 activity during DNA replication.

### MSH2 promotes G4 formation and limits PARP1 activity during DNA replication

To further test the idea that MSH2 limits G4 processing and in turn PARP1 replication activity, we analyzed the MSH2-null endometrial adenocarcinoma cell line, HEC59 in which chromosome 2 introduction establishes MSH2 expression^35^. Immunoblot revealed that the HEC59 (MSH2 deficient) and HEC59+chr2 (MSH2 proficient) cells display similar levels of total and CB-PARP1 (**Fig. 3a**). However, MSH2-restored HEC59+chr2 cells had more G4s along with reduced replication proficiency and RPA chromatin loading (**Fig. 3b-d**, **Supplementary Fig 3a, b**). Introduction of MSH2 also reduced PAR levels in both unchallenged conditions and following MMS and H_2_O_2_ treatment, again with PAR largely restricted to EdU positive cells (**Fig. 3e, f and Supplementary Fig. 3c**). Consistent with more PARP1 activity in the MSH2 deficient cells, sensitivity to PARPi was greater than the MSH2 proficient cells whereas as expected the MSH2 deficient cells were more resistant to alkylating agent, N-methyl-N’-nitro-N-nitrosoguanidine (MNNG)^35^ (**Fig. 3g, h**). Similar findings were observed in HeLa cells in which MSH2 KO maintained elevated RPA but reduced G4s (**Supplementary Fig. 3d-f**). PAR levels were also enhanced, along with PARPi sensitivity as compared to MSH2 proficient controls that had as expected greater MNNG sensitivity (**Supplementary Fig. 3g-i**). Together, these findings demonstrate that MSH2 aligns with not only enriched G4s but also reduced PARP1 S phase activity and targetability.

**Fig. 3:**
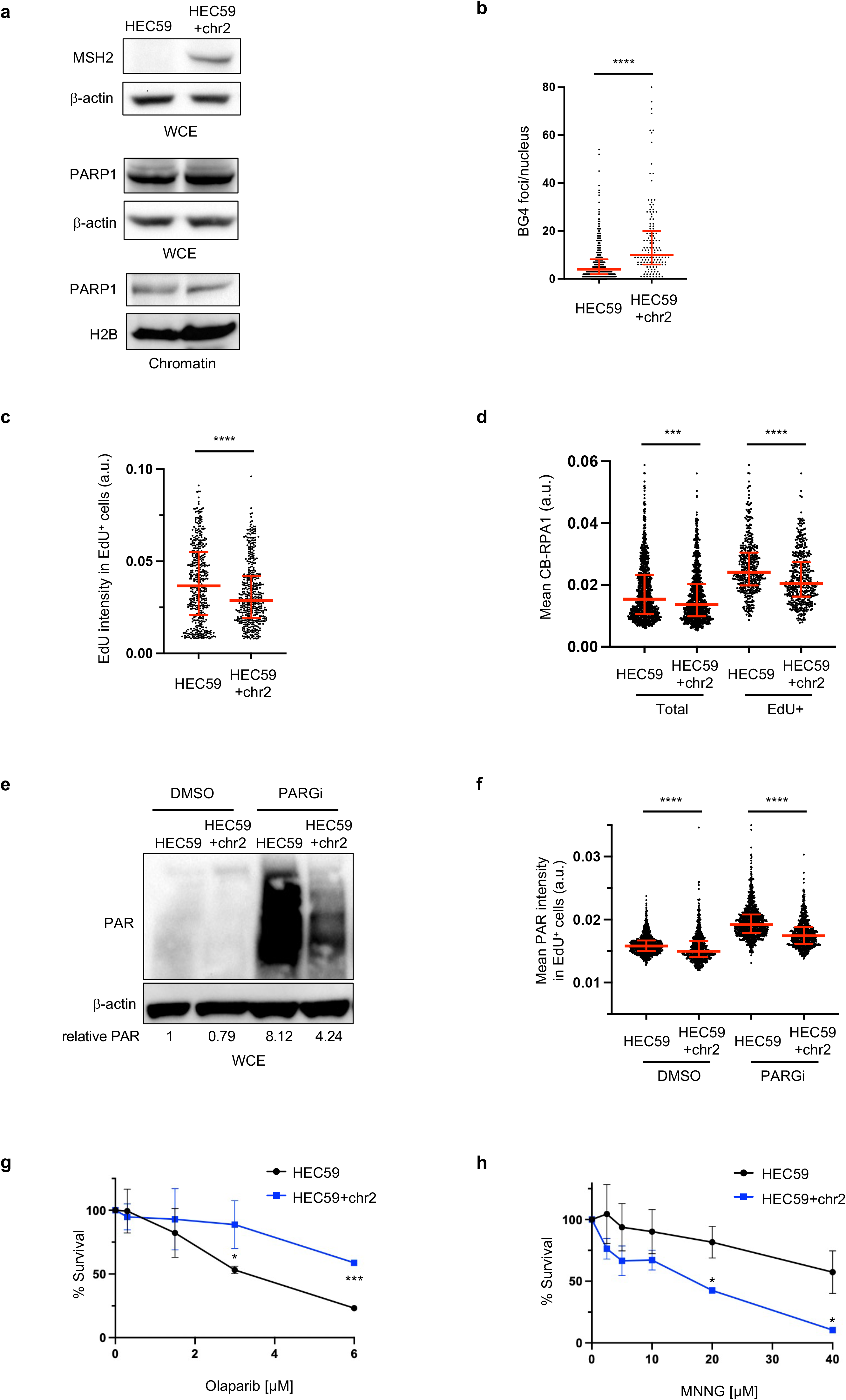
MSH2 interferes with PARP1 activation. **a** Representative WB experiment for whole cell lysates and chromatin fractionations showing indicated proteins in HEC59 vs HEC59+chr2 cells under untreated growth conditions. **b** Quantification of G4 in untreated HEC59 vs HEC59+chr2 cells. At least 100 cells are quantified from 3 biological independent experiments (n = 3). Red bars represent the median ± interquartile range. Statistical analysis according to Mann-Whitney test. **c** Quantification of mean EdU intensity from EdU^+^ (40min) cells. **d** Quantification of chromatin bound RPA1 (CB-RPA1) for the indicated cells with EdU incubated for 40 min. **e** Representative WB for the PAR formation in indicated cells treated with DMSO or PARGi (10 µM) for 40 min prior to harvesting. Quantification of mean relative PAR from three biological replicates normalized to β-actin shown. **f** Quantification of PAR after 30min DMSO or PARGi (10 µM) treatment in the indicated EdU^+^ cells. For the quantification in **c**, **d** and **f**, EdU^+^ cells were gated according to positive EdU incorporation. Each dot represents one cell. At least 300 cells are quantified from n = 3. Red bars represent the median ± interquartile range. All statistical analysis according to Kruskal-Wallis test, followed by Dunn’s test. **g, h** Cell survival assays for the indicated cells under increasing concentrations of Olaparib and MNNG. Data represent the mean percentage ± SD of survival for each dot. Significance was determined by unpaired t-test (two-tailed, unequal variance).

### MSH2 depletion restores PARPi sensitivity and gaps in cells lacking the FANCJ-MLH1 interaction

We next sought to evaluate if G4s and reduced PARP1 activity in cells lacking the FANCJ-MLH1 interaction was caused by MSH2. We first confirmed our previous finding that MSH2 depletion restored replication restart following release from APH or HU in cells lacking the FANCJ-MLH1 interaction (**Fig. 4a and Supplementary Fig. 4a-c**)^25^. Moreover, G4s were reduced (**Fig. 4b and Supplementary Fig. 4d**) and S phase PAR was elevated in both the RPE1 and FA-J cell systems (**Fig. 4c and Supplementary Fig. 4e, f**). We also observed that CB-PARP1 was reduced by MSH2 depletion and accordingly, we observed that PARPi led to a significant re-trapping of PARP1 (**Fig. 4d**). Finally, we observed that MSH2 depletion enhanced PARPi-induced replication speed as detected by the lengthening of DNA fibers (**Fig. 4e**) that can be toxic if discontinuous with gaps^17^. Indeed, we detected replication gaps^17,36^ by the presence of RPA loading and S1 nuclease sensitive DNA fibers (**Fig. 4e and Supplementary Fig. 4g**). Thus, in cells without the FANCJ-MLH1 interaction, MSH2 limits the capability of PARPi to promote speed and gaps indicating that PARP1 activation may be deficient. Indeed, MSH2 depletion enhanced PARPi sensitivity (**Fig. 4f and Supplementary Fig. 4h, i**), consistent with restored PARP1 catalytic activity. Notably, FANCJ null cells have a more muted response to MSH2 depletion (**Supplementary Fig. 4i**) as found with DNA interstrand crosslinking agents^25^, suggesting that this enhanced sensitivity requires FANCJ. Thus, it appears that MSH2 depletion restores FANCJ function in the FANCJ-MLH1 mutant cells consistent with MSH2 and FANCJ having opposing activities that are regulated by the FANCJ-MLH1 interaction.

**Fig. 4:**
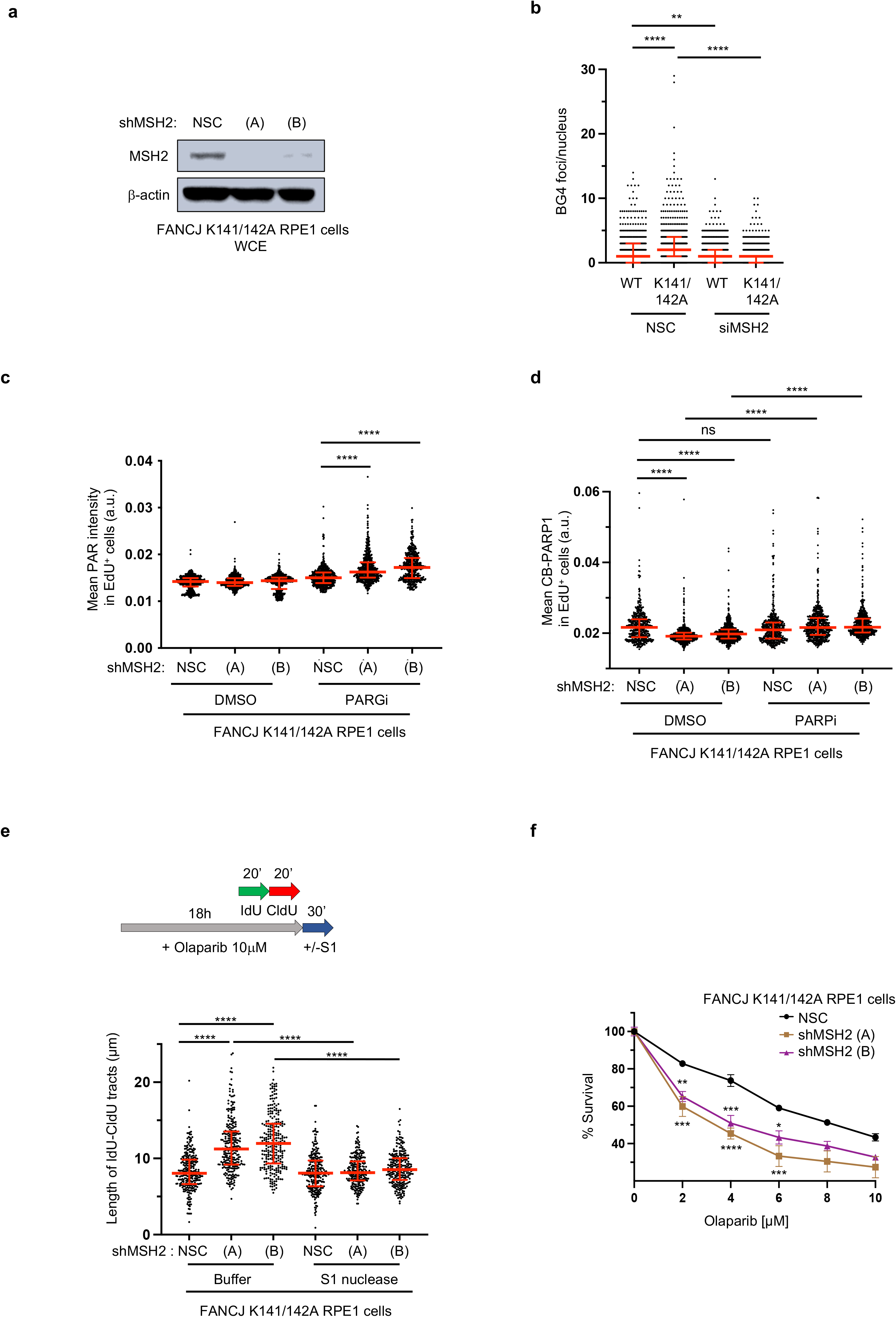
The FANCJ-MLH1 interaction promotes PARP1 activation by restricting MSH2. **a** Representative WB analysis with the indicated antibodies of extracts from RPE1 cells expressing small hairpin RNA (shRNA) against non-silencing control (NSC), MSH2 (A), and MSH2 (B). **b** Quantification of G4 in indicated RPE1 cells with siRNA under untreated growth conditions. **c** Quantification of PAR after 30min DMSO or PARGi (10 µM) treatment in the indicated EdU^+^ RPE1 cells. **d** Quantification of CB-PARP1 in the indicated cells with or without Olaparib treatment (10 µM, 6hrs), with EdU incubated at the final 40min. For the quantification from **b** to **d**, EdU^+^ cells were gated according to positive EdU incorporation. Each dot represents one cell. At least 300 cells are quantified from 3 biological independent experiments (n = 3). Red bars represent the median ± interquartile range. **e** Schematic and quantification of the S1 nuclease DNA fiber assays for the length of dual-color tracts in indicated cells following Olaparib treatment (10 µM, 18h). Each dot represents 1 fiber; at least 250 fibers are quantified from n = 3. Red bars represent the median ± interquartile range. **f** Cell survival assays for indicated RPE1 cells under increasing concentrations of Olaparib. Data represent the mean percentage ± SD of survival for each dot. Significance was determined by one-way ANOVA followed by Dunnett’s test comparing NSC to shMSH2 cells. For **b**, **c**, **d** and **e**, statistical analysis according to Kruskal-Wallis test, followed by Dunn’s test.

### G4 “sequestered” and canonical “trapped” PARP1 are distinct and provide insight into PARPi toxicity in BRCA deficient cells

FANCJ loss limits PARP1 activity which along with PARP1 trapping is thought to underlie the toxicity of PARPi. Thus, we sought to understand if PARP1 bound to G4s^28^ “G4-sequestered PARP1” was similar to canonical trapped PARP1 induced by PARPi (**Fig. 5a**). Canonical PARP1 trapping is detrimental in the context of XRCC1 null cells^37^ as validated by the finding that XRCC1 null cells are hypersensitive to PARPi yet insensitive to loss of PARP1 catalytic activity by PARP1 deletion (**Fig 5b, c**, **Supplementary Fig. 5a, b**), as reported^38,39^. If FANCJ depletion enhanced canonical trapped PARP1, FANCJ depletion could also sensitize XRCC1 null cells. However, FANCJ depletion did not reduce the colony formation of the XRCC1 null cells and instead modestly elevated PARPi resistance (**Fig. 5b, c**). Moreover, FANCJ deficient cells did not gain fitness following PARP1 deletion (**Fig. 5d**) suggesting that G4-sequestered PARP1 was not inherently toxic. However, as expected, BRCA1 depletion in PARP1 KO cells reduced clonogenic efficiency similar to the sensitivity detected with PARPi treatment (**Fig. 5d**) consistent with the dependency of BRCA1 deficient cells on PARP1^2,40–43^. Functional depletion of FANCJ or BRCA1 was confirmed immunoblot and by the sensitivity to PARPi versus cisplatin, respectively (**Supplementary Fig. 5c, d**). Overall, these findings indicated that FANCJ depletion creates a form of chromatin bound PARP1 that is not inherently toxic at least in FANCJ deficient or XRCC1 null cells.

**Fig. 5:**
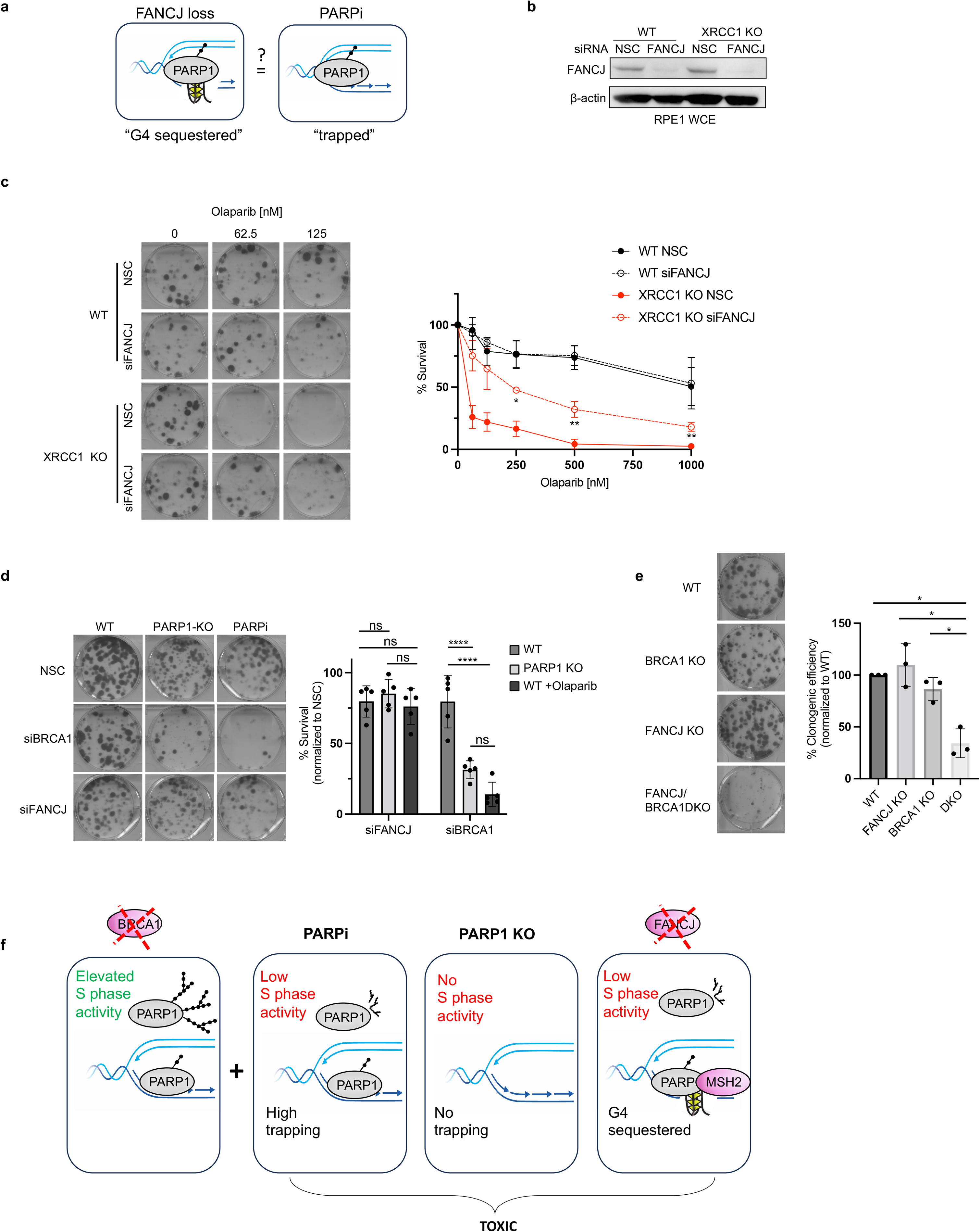
G4s “sequestered” PARP1 is distinct from “trapped” PARP1 and provides insight that loss of S phase activity underlies PARPi toxicity in BRCA1 deficient cells. **a** Model depicting the question, Is PARP1 trapping distinct from G4 sequestering? **b** Representative WB of whole cell lysates in indicated cells lines and knock down reagents. **c** Representative images and quantification of clonogenic survival assays of the indicated RPE1 cell lines and siRNA treatments with increasing doses of Olaparib, n=3 ± SD. Significance was determined by unpaired t-test (two-tailed, unequal variance) comparing XRCC1 KO NSC to siFANCJ. **d** Representative images and quantification of clonogenic assays for RPE1 WT and PARP1 KO cells with indicated siRNA, KO cell or Olaparib treatment (1 µM). Mean survival percentages normalized to non-silencing control (NSC), n=5 ± SD. Significance determined by two-way ANOVA followed by Tukey’s test. **e** Representative images and quantification of clonogenic assays for the indicated cells. DKO = double knockout. Mean percent clonogenic efficiencies (normalized to WT, n=3) ± SD. Significance determined by one-way ANOVA followed by Dunnett’s test. **f** Model of proposed findings: PARPi toxicity in BRCA1 deficient cells derives from an inability of PARP1 to function in S-phase resulting in the accumulation of replication-associated ssDNA lesions and PARP1 trapping. Loss of FANCJ in BRCA1 deficient cells sequesters PARP1 on G4s prohibiting PARP1 from efficiently functioning during replication.

Given that PARP1 binds G4s^28^ and is distinct from trapped PARP1, we sought to determine if this insight could be leveraged to clarify how PARPi is toxic in BRCA1 deficient cells. Indeed, PARP1 sequestering that limits PARP1 replication activity could be toxic in settings in which this PARP1 activity is essential. A cell system that could be dependent on PARP1 S phase activity is BRCA1 deficient cells that display elevated PARP1 S phase activity^17^. Moreover, BRCA1 deficient cells display increased PARylation when subjected to MMS and/or MMS followed by H_2_O_2_ as compared to wildtype or XRCC1 null cells that exhaust this activity^37^ (**Supplementary Fig. 5e, f**) indicating that PARP1 is readily activated in BRCA1 deficient cells. BRCA1 and FANCJ DKO cells were viable, however colony plating revealed that fitness was significantly curtailed compared to either single deficiency (**Fig. 5e**, **Supplementary Fig. 5g**) as further confirmed by siRNA depletion of BRCA1 or FANCJ in the single KO lines compared to WT (**Supplementary Fig. 5h**). This finding was similar to the greater than expected loss of viability between FANCJ and BRCA1 dual deletion as compared to each single deletion^18^. Collectively, these findings suggest that loss of PARP1 replication activity due to PARPi, PARP1 deletion or FANCJ deletion is toxic to BRCA1 deficient cells (**Fig. 5f**) predicting that PARPi efficacy in BRCA mutant cancer derives from loss of PARP1 S phase activity and not PARP1 trapping.

## DISCUSSION

BRCA1 and FANCJ are known to be involved in DNA double-strand break repair by homologous recombination, fork protection, and Fanconi anemia pathways. Moreover, mutations in these genes increase the risk of developing hereditary breast and ovarian cancer^14–16,44^. Based on these common functions, it was expected that akin to BRCA1 deficient cells, FANCJ deficient cells would be robustly sensitive to PARP inhibitors (PARPi), but they are not^17^. The present study aimed to explore the distinct role of FANCJ in PARPi induced gaps and sensitivity and to gain insight towards the relationship between gaps and PARP1 function and/or trapping. We find that without FANCJ, PARP1 S phase activity is abnormally low because PARP1 is “sequestered” in chromatin which is distinct from canonical chromatin trapping of PARP1. Accordingly, FANCJ depletion can elevate PARPi resistance in cells with sensitivity to canonical trapped PARP1. However, similar to PARPi or PARP1 deletion, FANCJ deletion in BRCA1 deficient cells is toxic. These findings highlight that the common feature of toxicity in BRCA1 deficient cells is loss of PARP1 replication activity as opposed to PARP1 trapping. Given that BRCA1 deficient cells have elevated PARP1 replication activity, our findings support that loss of PARP1 S phase activity is lethal in BRCA1 deficient cells because they rely on this activity to mitigate replication problems.

This detailed elucidation of FANCJ’s role in PARP1 activity, MSH2 localization, and G4 stabilization provide insight into cellular replication processes that impact PARPi efficacy and gap induction (**Supplementary Fig. 5i**). FANCJ, although not required for PARP1 chromatin localization, is critical in maintaining proper PARP1 activity levels and MMR pathway regulation. FANCJ deficient cells, observed in various types including RPE1, 293T and U2OS, exhibited conspicuously low PARP1 S phase activity. FANCJ deficiency did not inherently prevent PARP1 activation in response to DNA damage, indicated by maintained PARP1 activity in cells treated with DNA damaging agents like MMS and H_2_O_2_. Instead, our findings reveal that a FANCJ-MLH1 interaction counters MSH2-bound G4s that sequester and restrict PARP1 S phase activation. Consistent with this model, FANCJ deficient cells have elevated G4s, that are substrates for PARP1 and MSH2^26–33^, and correspondingly greater proximity between G4s and PARP1. Moreover, MSH2-deficient cells display more G4s and PARP1 activity compared to MSH2 proficient cells, which showed reduced PARP1 S phase activity and targetability. Our previous studies have proposed that FANCJ restricts the MSH2-MSH6 complex since both MSH2 and MSH6 are elevated in the replisome of FANCJ null cells, and depletion of MSH2 or MSH6, but not MSH3, rescues defects associated with FANCJ deficiency^16,25^. Moreover, given that MSH2 depletion restores PARP1 activity in cells lacking the FANCJ-MLH1 interaction, but not in cells without FANCJ, G4 unfolding ultimately requires FANCJ and may be opposed by MSH2 unless coordinated with MLH1. The MMR pathway is present at the replisome^45^, however in FANCJ null cells, MSH2/MSH6 becomes further enriched ^16^, thus we propose that this complex is uniquely restricted by FANCJ.

While our understanding of how PARP1 is activated during an undisturbed S phase to participate in backup lagging strand synthesis remains limited, our findings suggest that G4 structures and MMR possess the potential to inhibit this activity, thereby conferring resistance to PARPi. Foreseeably PARP1’s auto-inhibitory domain is not released when PARP1 binds negatively charged G4s as it would be at a DNA break. Indeed, FANCJ unfolds proteins, and unfolding of the PARP1 helical region is associated with its catalytic activation^46–48^, leading to the possibility that FANCJ unfolds and activates PARP1. FANCJ may also create a G4 structure that is amenable to PARP1 activation. Prior to unfolding G4s, FANCJ stabilizes G4s, and G4 stabilization is associated with ssDNA gaps and PARP1 activation^23,49,50^. Moreover, while like MMR, PARP1 has affinity for G4s, only specific G4s activate PARP1^29–32^. Furthermore, FANCJ displacement of MSH2 appears fundamental for PARP1 replication function. Intriguingly, MMR impacts centromeric DNA replication by binding secondary structures that limit RPA and checkpoint induction^51^ consistent with MMR occluding ssDNA. It remains to be determined whether a MMR-FANCJ pathway engagement and S phase PAR levels can ultimately serve as a pharmacodynamic marker of PARPi response^52^. However, the dependence of BRCA1 deficient cancer cells on PARP1 S phase activity predicts that loss of this activity such as by targeting FANCJ will be a therapeutic option.

The finding that FANCJ null cells readily tolerate the enriched PARP1 in chromatin also provides insight that not all “trapping” is the same. Indeed, canonical trapping underlies the low fitness and sensitivity of XRCC1-deficient cells to MMS^37^. Consistent with this interpretation, we find that these cells gain fitness as well as resistance to PARPi upon by PARP1 loss. Given that XRCC1-deficient cells also gain PARPi resistance with FANCJ loss, we can surmise that canonical trapping is also reduced in this setting. Indeed, PARPi sensitivity in several genetic contexts is reversed by PARP1 deletion consistent with a trapping model^18,53–55^. Likewise, in the context of BRCA1 mutant cells in which residual BRCA function tolerates PARP1 loss, PARPi toxicity can be suppressed by mutations that reduce PARP1 trapping^56–58^. Moreover, cancer cells with distinct BRCA mutations, gain resistance by loss of PARP1^40,58^.

By contrast, when PARP1 loss causes a major loss to fitness, PARPi is likely toxic due to disruption of PARP1 S phase activity and PARP1 trapping may be less consequential. Indeed, the finding that FANCJ loss is toxic with BRCA1 loss, highlights that loss of PARP1 activity could be the fundamental issue. This possibility is supported by the finding that BRCA deficient cells are reported to be sensitive to either PARPi or PARP1 loss^2^. Our own findings also align with this conclusion. The idea that PARP1 activity in S phase maintains the homeostasis in BRCA deficient cells is furthered by the finding that PARP1 activity is elevated as shown previously and herein^17^. Furthermore, the role of BRCA proteins in preventing replication-associated gaps sheds new light on why PARP1 activity could be essential. Dual loss of BRCA and PARP1 and the combined replication dysfunction creates a toxic level of gaps that grossly limits cell fitness^13,17,59^. This phenomenon has also been observed with combined loss of polymerase theta (Pol θ) and BRCA^60–62^. The theory postulating that the toxicity of PARPi in BRCA deficient cells is attributed to the drug’s capacity to trap PARP1, substantiated by discoveries indicating that more potent trappers exhibit increased toxicity to these cells, has recently faced skepticism, spurred by emerging evidence suggesting that more potent trappers inherently possess enhanced catalytic inhibitory strength^10,63,64^.

In summary, our research sheds light on the complex mechanisms underlying PARP inhibitor sensitivity in different genetic contexts and highlights that in BRCA deficient cells, the anti-cancer mechanism is due to loss of PARP1 S phase activity. We believe that these findings have significant implications for the development of targeted therapies and precision medicine approaches for treating cancer. Furthermore, our findings highlight that depending on underlying mechanism of action, PARPi resistance will also evolve in distinct ways.

## ACKNOWLEDGEMENTS

We thank the members of the Cantor laboratory for helpful discussions. We thank Dr. Daniel Durocher for RPE1 cell lines including WT, BRCA1 K/O and isogenic matched RPE1 cells, Dr. Keith Caldecott for the XRCC1 KO and matched isogenic WT RPE1 cells, and Dr. Christopher Heinen for the HeLa MSH2 K/O and matched isogenic HeLa WT cells. This work was supported by R01 CA254037, R01 CA225018-02, Fanconi Anemia Research Fund, and charitable contributions from Mr. and Mrs. Edward T. Vitone, Jr (Cantor), and NIH K01 AG056554 and an NSF CAREER Award (Day).

## AUTHOR CONTRIBUTIONS

S.C, T.D., N.M, and K.C., designed the experiments. K.C., N.M., S.L., S.G.M., J.C., M.P., and A.N.K. performed the experiments. K.C., N.M., S.L., and J.C. analyzed the data. S.C., N.M, and K.C., wrote the manuscript. S.C., and T.D., supervised the research.

## METHODS

### Cell lines and related reagents

Human RPE1-hTERT (*TP53^−/−^*), 293T, U2OS and HeLa derived cell lines were grown in DMEM supplemented with 10% fetal bovine serum (FBS, Sigma-Aldrich), 1% penicillin/streptomycin (Gibco) and 1% Sodium pyruvate (only for RPE1). The generation of *BRCA1* KO and *FANCJ* KO RPE1-hTERT (*TP53^−/−^*), 293T and U2OS cell lines were described before^16,17^. The *BRCA1* and *FANCJ* double KO cells were generated similarly as before. *PARP1* and *XRCC1* related WT and KO RPE1 cells were from Caldecott lab^37^. Human endometrial HEC59 and HEC59+chr2 cell lines, FA-J (EUFA30-F) cell lines were cultured in DMEM supplemented with 15% FBS. For the RPE1-hTERT (*TP53^−/−^*) *FANCJ* gene knock-in cell lines, the desired *FANCJ* gene variants (K141/142A, K52R) were introduced in the RPE1-hTERT (*TP53^−/−^*) cells according to the RNP CRISPR approach of IDT. Detailed steps, sequences of PCR primers, sgRNA and ssODN repair templates can be found in table below. The expression of FANCJ variants were confirmed by immunoblot analysis.

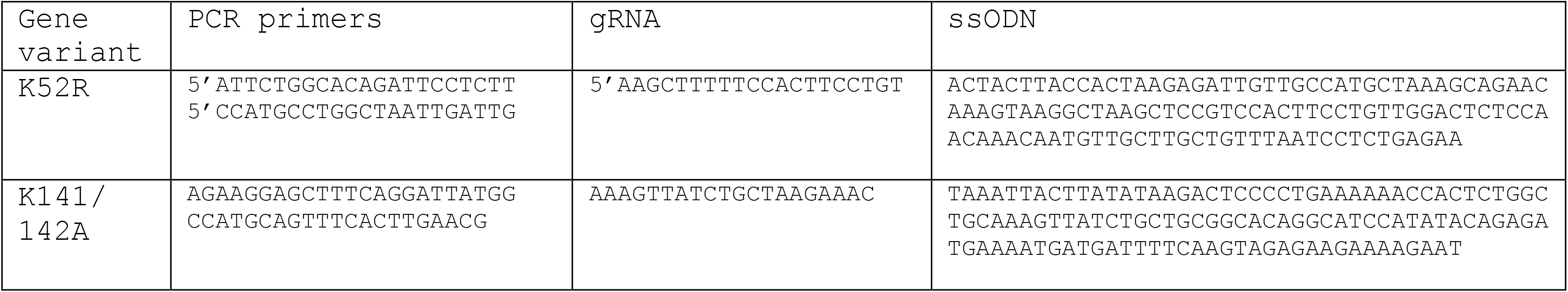

### Drugs and other reagents

The following drugs were used in the course of this study: PARP inhibitor Olaparib (SelleckChem AZD-2281 S1060). Cisplatin (Sigma-Aldrich P4394), camptothecin (CPT, Sigma-Aldrich C9911), Methyl methansulfonate (MMS, Sigma-Aldrich 129925), PARG inhibitor (PDD 0017273, Tocris 5952), Aphidicolin (Sigma A0781), Hydroxyurea (Sigma-Aldrich H8627), Hydrogen peroxide solution (H_2_O_2_, H1009, Sigma-Aldrich), Mitomycin C (MMC, MA4287, Sigma-Aldrich), Dimethyl sulfoxide (DMSO, Sigma-Aldrich D5879). All drugs were directly used or prepared per the manufacturer’s instructions. Other reagents used in this study included 5-chloro-2’-deoxyuridine (CldU, Sigma-Aldrich C6891), 5-Iodo-2’-deoxyuridine (IdU, Sigma-Aldrich I7125). Concentration and duration of treatment are indicated in the corresponding figures and sections.

### Immunoblotting

Cells were harvested, lysed, and processed for western blot analysis as described previously using 150 mmol/L NETN lysis buffer [20 mmol/L Tris; (pH 8.0), 150 mmol/L NaCl, 1 mmol/L EDTA, 0.5% NP-40, and Halt Protease inhibitor cocktail (Thermo Fisher Scientific 78440)]^17^. For cell fractionation, cytoplasmic and soluble nuclear fractions were isolated with the NE-PER Kit (Thermo Fisher Scientific 78835) according to the manufacturer’s protocol; to isolate the chromatin fraction, the insoluble pellet was resuspended in RIPA buffer (Cold Spring Harbor Protocol) and followed by 15min sonication by Diagenode bioruptor with medium power for 30s on and 30s off at 4 °C. Proteins were separated using SDS–PAGE and electro-transferred to nitrocellulose or polyvinylidene difluoride membranes. Membranes were blocked in 5% non-fat dry milk (NFDM) in Tris-buffered saline with 0.1% Tween-20 and incubated with primary antibody overnight at 4 °C. Primary antibodies for western blot analysis included anti-FANCJ (E67 from Cantor lab), anti-β-actin (Sigma-Aldrich A1978), anti-PAR (poly-ADP-ribose binding reagent, Millipore Sigma MABE1031), anti-PARP1 (Abcam ab227244), anti-H2B (Cell Signaling Technology 8135), anti-MSH2 (Abcam ab52266), anti-XRCC1 (Novus Biological NB120-1838). Secondary antibodies include ECL anti-rabbit IgG, HRP-linked whole antibody (from donkey, GE Healthcare NA934) and ECL anti-mouse IgG, HRP-linked F(ab’)_2_ fragment (from sheep, Thermo Fisher Scientific NA9310) All antibodies were used within the range of suggested dilution. Membranes were washed, incubated with corresponding horseradish peroxidase-linked secondary antibodies (Amersham, GE Healthcare) for 1hour at room temperature (RT) and detected by chemiluminescence imaging system (Bio-Rad) or Kodak X-OMAT 2000A Film Processor. Quantification of immunoblots were done with ImageJ by normalizing PAR to β-actin and calculating relative PAR levels with respect to the WT untreated sample.

### Plasmids and RNA interference

Lentiviral production was described previously in detail^65^. Similarly, the mutant clones of FANCJ K141/142A and K52R were individually generated using site-directed mutagenesis (Genscript) with PMT-BRD025 FANCJ WT (wild-type) as the template, and were sequence verified (Genscript Piscataway). Details of standard virus production pipelines can be found at the Broad Institute’s Genetic Perturbation Platform website (https://portals.broadinstitute.org/gpp/public/). Viruses for the mutant and WT FANCJ were produced in 96-well plates using HEK293T cells transfected with packaging vector psPAX2 (100 ng), envelope plasmid CMV-VSVG (10 ng), and respective PMT-BRD025 FANCJ mutant plasmid (100 ng). Lentiviral-containing supernatants were harvested 31 hours post-transfection and stored in polypropylene plates at −80 °C until use.

Stably transduced cells were generated by infection with pLKO.1 vectors containing shRNAs against non-silencing control (NSC) or one of the shRNAs against corresponding genes: MSH2 includes (A) 5’-AGCAAGCTCTGCAACATGAAT-3’, (B) 5’-TTACCTTCATTCCATTACTGG-3’. The information was obtained from Dharmacon website (https://horizondiscovery.com), and the shRNAs were obtained from the University of Massachusetts Chan Medical School shRNA core facility. Cells were selected by puromycin for 3-5 days before experiments were carried out. Transfection of siRNA in RPE1 cells was performed with Lipofectamine 3000 (Thermofisher) according to the manufacturer’s instructions. Briefly, cells were plated in 8-well slides and each well was transfected with 1 μl of Lipofectamine 3000 and contained a final siRNA concentration of 36 nM in a total volume of 0.5 mL. Assays were performed at 48 hr post transfection. siRNA used were ON-TARGETplus Human MSH2 (4436) SMARTpool (L-003909) and ON-TARGETplus Non-targeting Pool (D-001810). For studies involving depletion of BRCA1 or FANCJ, cells were plated in 6-well dishes. The next day, each well received 3 µL of DharmaFECT 1 Transfection Reagent (Horizon Discovery) and a final siRNA concentration of 25 nM according to the manufacturer’s instructions. Dharmacon siRNA used were siLuciferase (D-002050-01-20 or D-001210-02-50), siBRCA1 SMARTpool (M-003461-00-05), and siFANCJ (5′ GUACAGUACCCCACCUUAU 3′).

### Viability assays

Cells were seeded onto 96-well plates (300 cells per well, performed in biological triplicates for each experiment group) and incubated overnight. The next day, cells were treated with increasing doses of drugs as indicated in corresponding figures and maintained in complete media for 5 to 7 days. Percentage survival was measured photometrically using CellTiter-Glo 2.0 (Promega G9242) viability assay in a microplate reader (Beckman Coulter DTX 880 Multimode Detector). For clonogenic survival assays, 250-500 cells per well were seeded into 6-well plates. The next day, the media was replaced with media containing treatments or vehicle and incubated for 10 to 14 days. Percentage survival was determined by manual cell counting of at least 50 cells or more^66^ after staining with crystal violet staining solution (0.05% w/v crystal violet, 1% formaldehyde, 1% MeOH, in 1x PBS) (Sigma-Aldrich C0775).

### Immunofluorescence

For Poly(ADP-ribose) or PAR, cells cultured on coverslips were fixed with 4% formaldehyde in PBS for 10 min at RT and subsequently permeabilized by a 5 min incubation in ice-cold methanol/acetone solution (1:1). After blocking the cells with 10% fetal calf serum for 30 min (alternatively add 3% BSA), coverslips were incubated with the primary antibody anti-PAR polyclonal antibody (Trevigen 4336-BPC-100) at 37 °C for 1hr. Followed by PBS washing, cells were then incubated with the appropriate fluorescently labeled secondary antibody for 1hr at RT. EdU labeling was performed using Click-iT EdU Alexa Fluor 488 Imaging Kit (Invitrogen C10337) according to the manufacturer’s instructions. Coverslips were then washed, stained with DAPI (1 mg/ml in PBS, Thermo Fisher Scientific D1306) for 30 min and mounted using VECTASHIELD mounting media (Vector Laboratories H-1200).

For chromatin bound proteins, cells on coverslips were plated on ice for 0.5-1 min before pre-extracted by ice-cold PBS+0.5% Triton for 5 min. Then, cells were fixed by 3%paraformaldehyde/2%sucrose for 10 minutes at RT. Cells were washed twice with PBS-T (0.01% Tween) and incubated with primary antibodies (anti-PARP1 antibody, Abcam ab227244; anti-MSH2 antibody, Abcam ab52266; anti-RPA1 antibody, Cell Signaling Technology 2267) in DMEM + 10% FBS (alternatively add 3% BSA) at 37C for 1 hr. After 3x PBS-T washing, coverslips were incubated with appropriate secondary antibodies (Alexa Fluor 488 goat anti-Mouse A-11001 and Alexa Fluor 568 goat anti-Rabbit A-11011) in DMEM + 10% FBS and DAPI. EdU labeling was performed as described above. Finally, after washing with PBS-T (x3), coverslips were mounted with Prolong (Invitrogen P36930). For all assays above, images were collected by fluorescence microscopy (Axioplan 2 imaging and Axio Observer, Zeiss) at a constant exposure time in each experiment. Representative images were processed by ImageJ software. Mean intensity of immunofluorescence for each nucleus were measured with Cell Profiler software version 3.1.5 from Broad Institute.

For BG4 Immunofluorescence, RPE1 and HEC59 cells were cultured overnight in 8-well slides (Nunc Lab-Tek II Chamber Slide) with 50,000 cells per well. Cells were washed in PBS and then fixed in 4% paraformaldehyde for 10min at room temperature. Following three PBS washes, cells were permeabilized with 0.1% Triton-X in PBS. Cells were treated with 0.24mg/ml Monarch RNase A (New England Biolabs) for 1 hour at 37°C. After a PBS wash, cells were blocked with 0.5% goat serum, or alternatively 0.5% FBS, in PBS for 1 hour at 37°C, followed by incubation with BG4 antibody at a 1:100 dilution in 0.1% Tween20 in PBS (PBST) and 0.5% goat serum at 37°C for 1 hour. The BG4 antibody was prepared according to^67^, DYKDDDDK Anti-FLAG antibody (cat. no. 14793S, Cell Signaling Technology) was used at 1:800 in 0.1% PBST and 0.5% goat serum for 1 hour at 37 °C. Following three washes in 0.1% PBST, cells were incubated with 1:1000 Alexa Fluor 594 goat anti-rabbit secondary antibody (cat. no. A-11012, Life Technologies) in 0.1% PBST and 0.5% goat serum for 1 hour at room temperature. Finally, cells were overlayed with VECTASHIELD Antifade Mounting Media with DAPI (cat. no. H-1200-10, Vector Laboratories) before sealing. Fluorescent foci were imaged using a Zeiss LSM 710 confocal microscope (objective magnification: 40x, oil). Each experiment was performed with three biological replicates. N=5 images were analyzed for each condition using a custom ImageJ 2.9.0 script and GraphPad Prism 9.4.1. Where mean fluorescence intensity is reported,images were taken at 63x with fluorescence microscopy (Axioplan 2 imaging and Axio Observer, Zeiss) and analyzed with Cell Profiler software version 4.2.1 from the Broad Institute.

### Proximity ligation assay

Cells were seeded on 18 mm x 18 mm coverslips and the following day they were pulsed with 10mM EdU for 20’. After the EdU pulse, cells were initially pre-fixated with 0.1% formaldehyde in PBS for 2’ at RT. Following three PBS washes, the samples were pre-extracted with CSK buffer on ice for 5’ and then washed with PBS before fixing with 3.7% formaldehyde at RT for 15’. Coverslips were then washed with PBS and stored O/N at 4 °C. The following day cells were post-fixated and permeabilized with methanol for 20’ at −20C. The samples were washed 3X with PBS and blocked for 30’ with 3% BSA in PBS. To label EdU, a click reaction (100mM Tris pH 8, 100mM CuSO4, 2mg/ml sodium-L-ascorbate, 10mM biotin-azide) with Alexaflour 488 azide was performed for 30’. Slides were then incubated with primary antibodies for 1 hour at 37 °C (1:250 mouse anti-MSH2 Abcam ab52266; 1:500 rabbit anti-PARP1 Abcam ab227244; 1:500 rabbit anti-PCNA Abcam ab18197, 1:500 mouse anti-PCNA Abcam ab29 diluted in blocking solution). After antibody incubation, coverslips were washed 2X with Buffer A for 5’at RT (Duolink kit DUO92101). Each coverslip was then incubated for 1 hour at 37 °C with Duolink PLA probes (Thermo Fisher Scientific) diluted in blocking solution. After 2X washes with Buffer A for 5’at RT, probes were ligated for 30’ at 37 °C and amplified by polymerase reaction for 100’ at 37 °C. Coverslips were then washed 2X with Buffer B for 5’at RT (Duolink kit) and then mounted with DAPI on microscope slides. Images were acquired by fluorescence microscopy (Axioplan 2 imaging and Axio Observer, Zeiss) with a 63X objective. Deconvolution of the images was done using the ImageJ software. The number of foci in each cell was counted with Cell Profiler software and the statistical analysis was performed using Prism (GraphPad Software).

For the BG4-PARP1 PLA, the above protocol was modified to include the following steps. After post-fixation and permeabilization with methanol for 20’ at −20C, the samples were washed three times for 5’ each with PBS-0.01%Tween and incubated with 0.24 mg/ml Monarch RNase A (New England Biolabs) for 1 hour at 37°C. After three PBS washes, the cells were blocked with PBS-0.1%Tween and 10% FBS for 30’ at 37 °C. Next, the BG4 antibody^67^ was incubated for 1 hour at 37°C and subsequently incubated with 1:800 mouse anti-FLAG antibody (F1804, Sigma) to BG4 and 1:500 rabbit anti-PARP1 (ab227244, Abcam) for 1 hour at 37°C in PBS-0.1%Tween and 10% FBS. The standard PLA protocol described above was followed to complete sample processing.

### DNA fiber assay

Similar as previously described^17^, cells were labeled by sequential incorporation of two different nucleoside analogs, IdU and CldU, into nascent DNA strands for the indicated time and conditions. After nucleoside analogs were incorporated in vivo, the cells were collected, washed, spotted, and lysed on positively charged microscope slides by 7.5 mL spreading buffer for 8 min at room temperature. For experiments with the ssDNA-specific endonuclease S1, after the CldU pulse, cells were treated with CSK100 buffer for 10 min at room temperature, then incubated with S1 nuclease buffer with or without 20 U/mL S1 nuclease (Invitrogen, 18001-016) for 30 min at 37 °C. The cells were then scraped in PBS + 0.1% BSA and centrifuged at 7,000 rpm for 5 min at 4 C. Cell pellets were resuspended at 1,500 cells/mL and lysed with lysis solution on slides. Individual DNA fibers were released and spread by tilting the slides at 45 degrees. After air-drying, fibers were fixed by 3:1 methanol/acetic acid at room temperature for 3 min. After air-drying again, fibers were rehydrated in PBS, denatured with 2.5 M HCl for 30 min, washed with PBS, and blocked with blocking buffer (PBS + 0.1% Triton + 3%BSA) for 1 hr. Next, slides were incubated for 2.5 hr with primary antibodies for (IdU, Becton Dickinson 347580; CldU, Abcam 6326) diluted in blocking buffer, washed several times in PBS, and then incubated with secondary antibodies (IdU, goat anti-mouse, Alexa Fluor 488; CldU, goat anti-rat, Alexa Fluor 594) in blocking buffer for 1 hr. After washing and air-drying, slides were mounted with Prolong (Invitrogen P36930). Finally, green and/or red signals (measure at least 100 fibers for each experiment) were visualized by fluorescence microscopy (Axioplan 2 imaging, Zeiss) for the active replication at the single-molecule level.

### Statistical analysis

Statistical differences in DNA fiber assays and immunofluorescence intensity were determined by nonparametric Kruskal-Wallis test followed by Dunn’s test for multiple comparations in non-Gaussian populations. Two group comparations were determined using two-tailed Mann-Whitney test. Statistical differences in viability assays with small sample sizes were determined by two-way ANOVAs followed by Tukey’s test (two-independent variables with multiple comparisons), one-way ANOVAs followed by Dunnett’s test (multiple comparisons) or unpaired t-test (two-group comparison, two-tailed, unequal variance). Statistical analysis was performed using GraphPad Prism (Version 9.0). In all cases, ns: not significant (p > 0.05), *p < 0.05, **p < 0.01, ***p < 0.001, and ****p < 0.0001.

### Data availability

Unprocessed blots, gels, and microscopy images are available online and through lead contact upon request. All materials associated with this study are available upon request from the lead contact. Further information and requests for resources and reagents should be directed to and will be fulfilled by the Lead Contact, Sharon Cantor.

## FIGURE TITLES AND LEGENDS

**Supplementary Figure 1:**
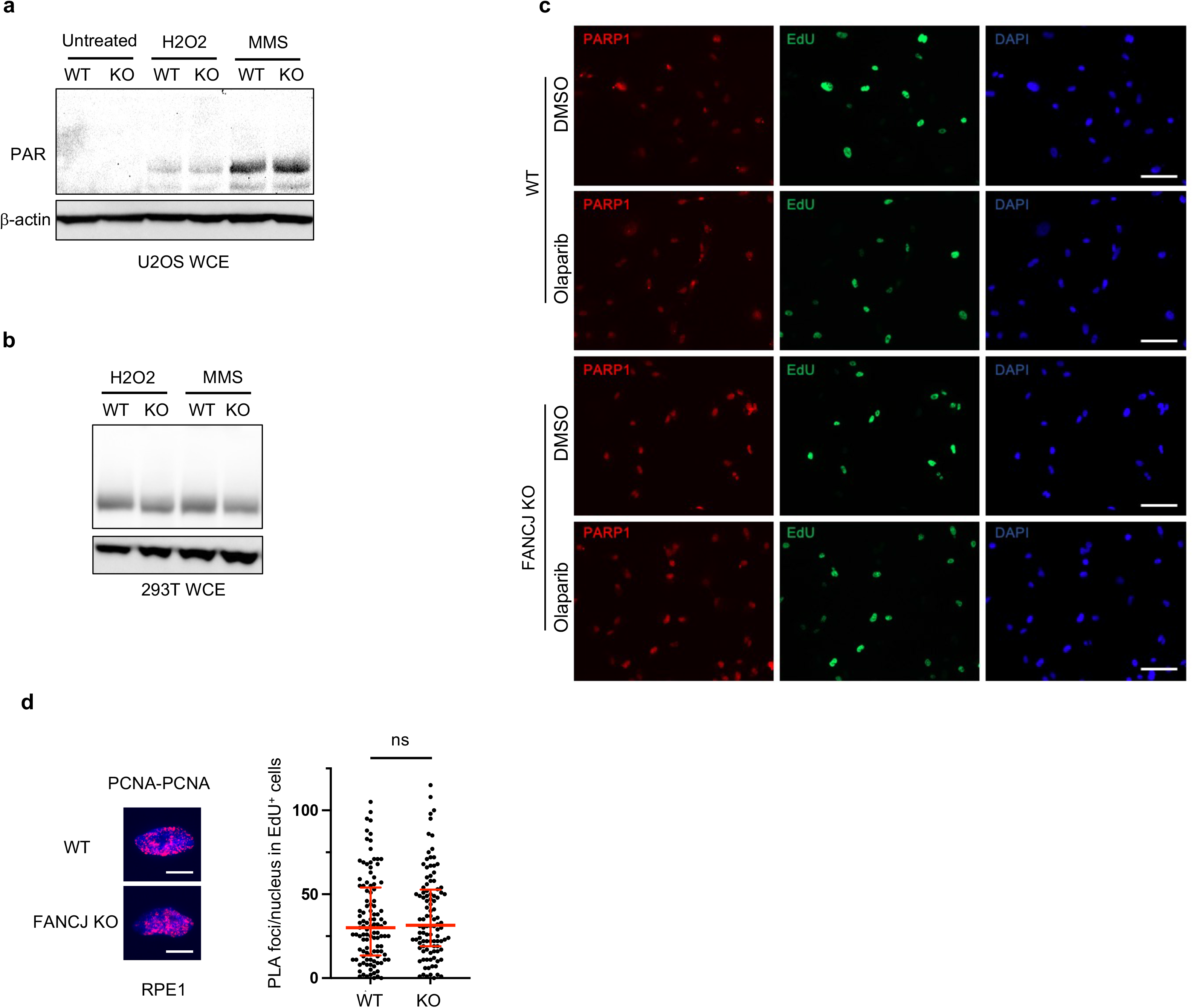
FANCJ deficiency does not reduce PAR response to DNA damage. **a, b** Representative WB for PAR in indicated cells under untreated growth conditions or following the treatment of DNA damage-inducing agents methyl methanesulfonate (MMS) and H_2_O_2_ for 40 min. **c** Representative images related to Fig 1h. Scale bars, 50 µm **d** PCNA-PCNA PLA assay in untreated RPE1 WT vs FANCJ KO cells. Dot plot shows the number of foci and the median ± interquartile for at least 100 cells from 3 biological independent replicates (n=3). Scale bars, 10 µm. Statistical analysis according to Mann-Whitney test.

**Supplementary Figure 2:**
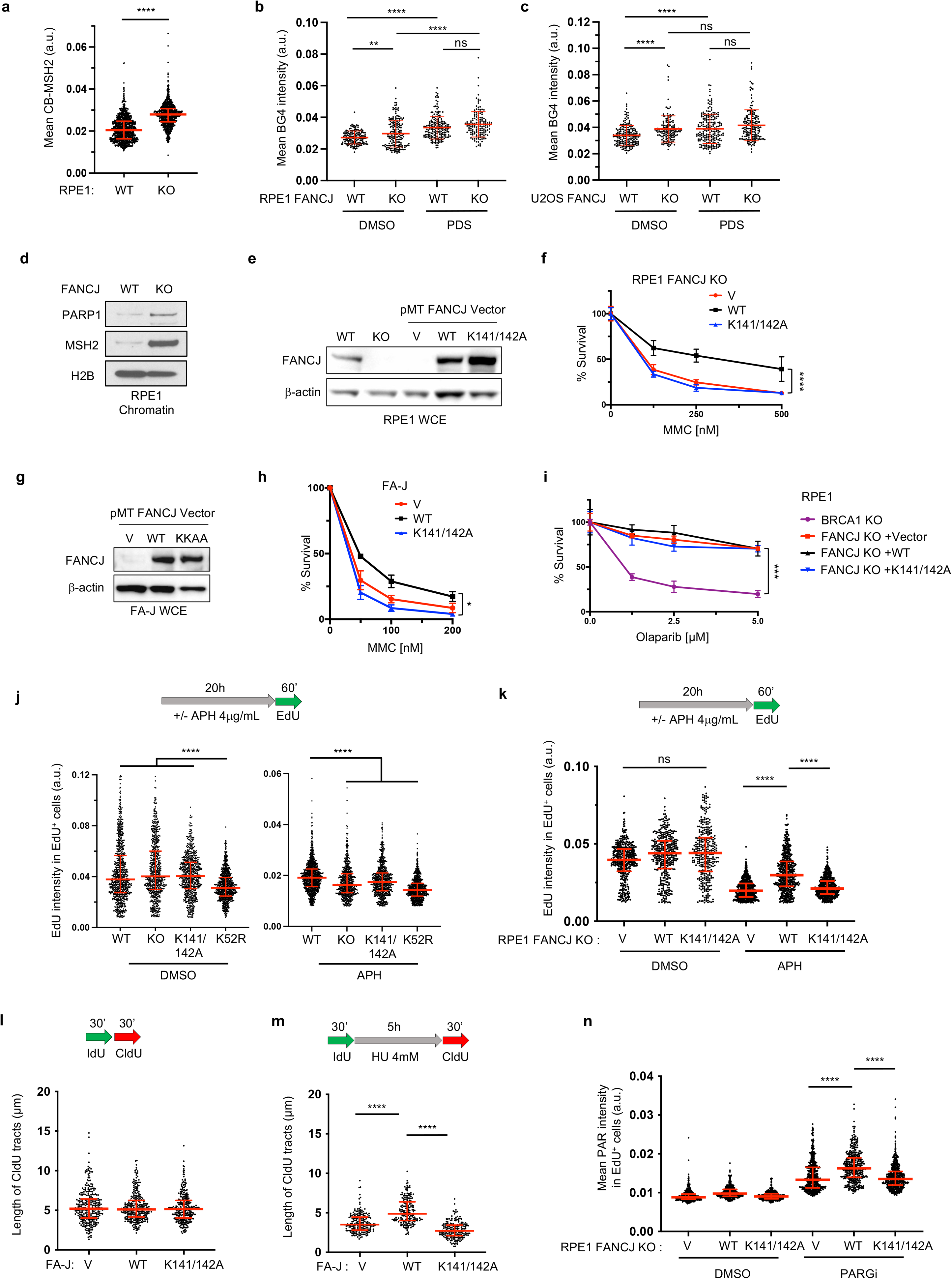
Loss of FANCJ or FANCJ-MLH1 binding impairs RPA chromatin loading and replication restart. **a** Quantification of chromatin-bound MSH2 (CB-MSH2) for indicated RPE1 cells. Statistical analysis according to Mann-Whitney test. Each dot represents one cell, at least 200 cells are quantified from n = 2. Statistical analysis according to Mann-Whitney test. **b, c** Mean BG4 fluorescence intensity in RPE1 and U2OS WT cells or FANCJ KO with or without 30’ 2 µM PDS. Each dot represents one cell, at least 150 cells are quantified from n = 2. Statistical analysis according to Kruskal-Wallis test followed by Dunn’s test. **d** Representative WB of chromatin bound PARP1 and MSH2 in RPE1 WT and FANCJ KO cells (n=2). **e** WB analysis for the whole cell lysates from FANCJ KO RPE1 cells complemented with FANCJ mutants. **f** Cell survival assays validating the complement system in RPE1 FANCJ KO cells. Data represent the mean percentage ± SD of survival for each dot. **g, h** WB analysis and cell survival assays validating the FANCJ mutant complemented in FA-J cells. **i** Cell survival assays for indicated cells (RPE1 complement system) under increasing concentrations of Olaparib. Data represent the mean percentage ± SD of survival for each dot. For **f**, **h** and **i**, significance was determined by one-way ANOVA followed by Dunnett’s test. **j, k** Quantification of mean EdU intensity from EdU^+^ cells. EdU was incubated for 1hr after 20 hr DMSO or aphidicolin (APH, 4 µg/ml) treatment in the indicated cells: FANCJ mutant cells **g** and FANCJ K141/142A cells in complement systems **h**. **l, m** Schematic and quantification of the DNA fiber assays for the length of IdU-connected CldU tracts in indicated cells following (**l**) without or (**m**) with HU treatment (4 mM, 5 h). Each dot represents 1 fiber; at least 200 fibers are quantified from n = 2. Red bars represent the median ± interquartile range. Statistical analysis according to Kruskal-Wallis test, followed by Dunn’s test. **n** Quantification of PAR intensity for FANCJ K141/142A cells in complement systems. For the quantification in **j**, **k**, and **n**, EdU^+^ cells were gated according to positive EdU incorporation. Each dot represents one cell. At least 200 cells are quantified from n = 2. Red bars represent the median ± interquartile range. All statistical analysis according to Kruskal-Wallis test, followed by Dunn’s test.

**Supplementary Figure 3:**
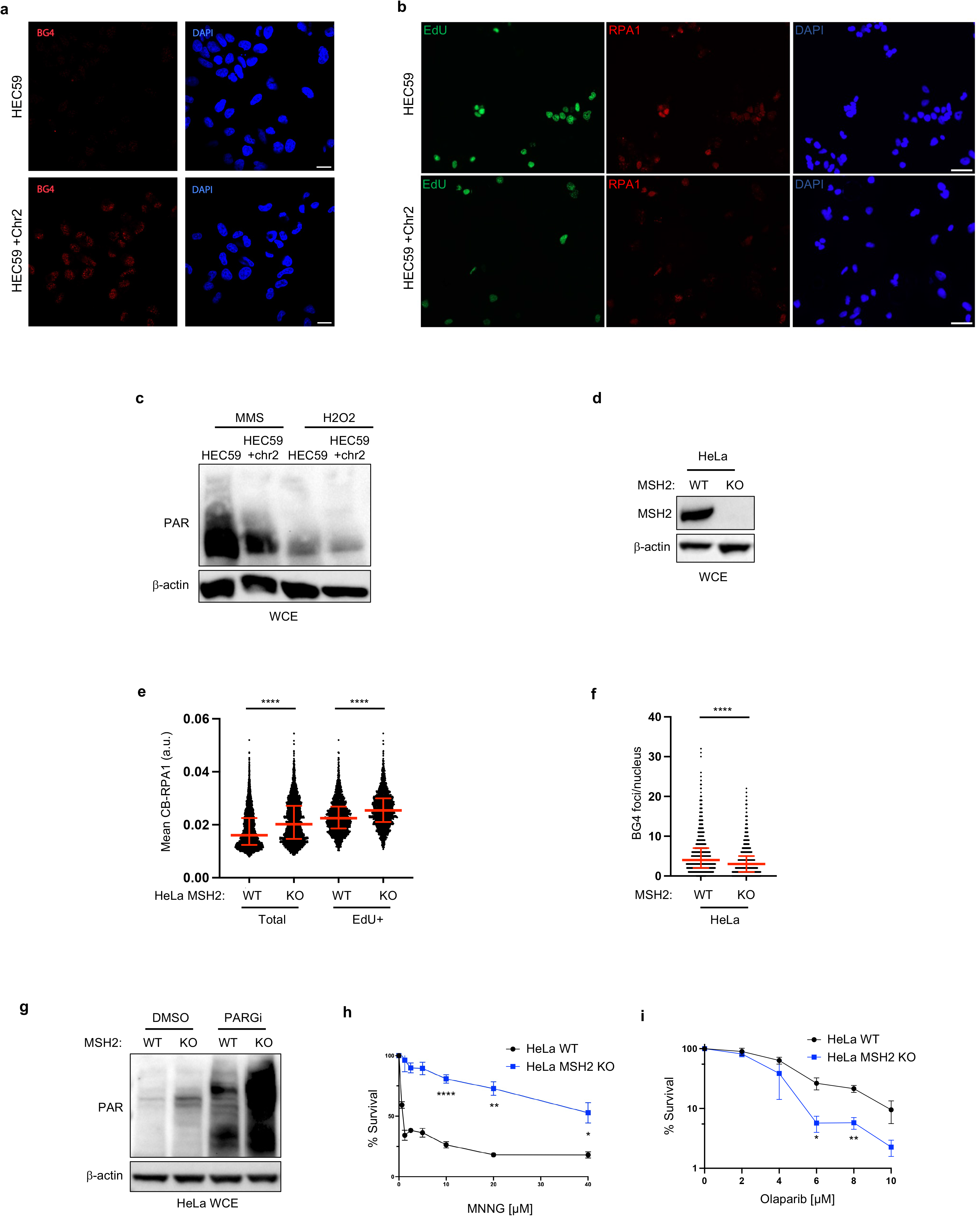
MSH2 interferes with PARP1 activation. **a** Representative images related to Fig 3b. Scale bars, 20 µm **b** Representative images related to Fig 3c, d. Scale bars, 50 µm **c** Representative WB for the PAR formation in indicated cells treated with MMS or H_2_O_2_ for 40 min prior to harvesting. **d** Representative WB experiments for the whole cell lysates showing indicated proteins in untreated HeLa WT vs HeLa MSH2 KO cells. **e** Quantification of chromatin bound RPA1 for the indicated cells with EdU incubated for the 40 min. EdU+ cells were gated according to positive EdU incorporation. Each dot represents one cell. At least 300 cells are quantified from n = 3. Red bars represent the median ± interquartile range. All statistical analysis according to Kruskal-Wallis test, followed by Dunn’s test. **f** Quantification of G4 in untreated HeLa WT vs HeLa MSH2 KO cells. At least 1000 cells are quantified from 3 biological independent experiments (n = 3). Red bars represent the median ± interquartile range. Statistical analysis according to Mann-Whitney test. **g** Representative WB for PAR in the indicated cells treated with DMSO or PARGi for 40 min prior to harvesting. **h, i** Cell survival assays of the indicated cells under increasing concentrations of MNNG and Olaparib. Data represent the mean percentage ± SD of survival for each dot. Significance was determined by unpaired t-test (two-tailed, unequal variance).

**Supplementary Figure 4:**
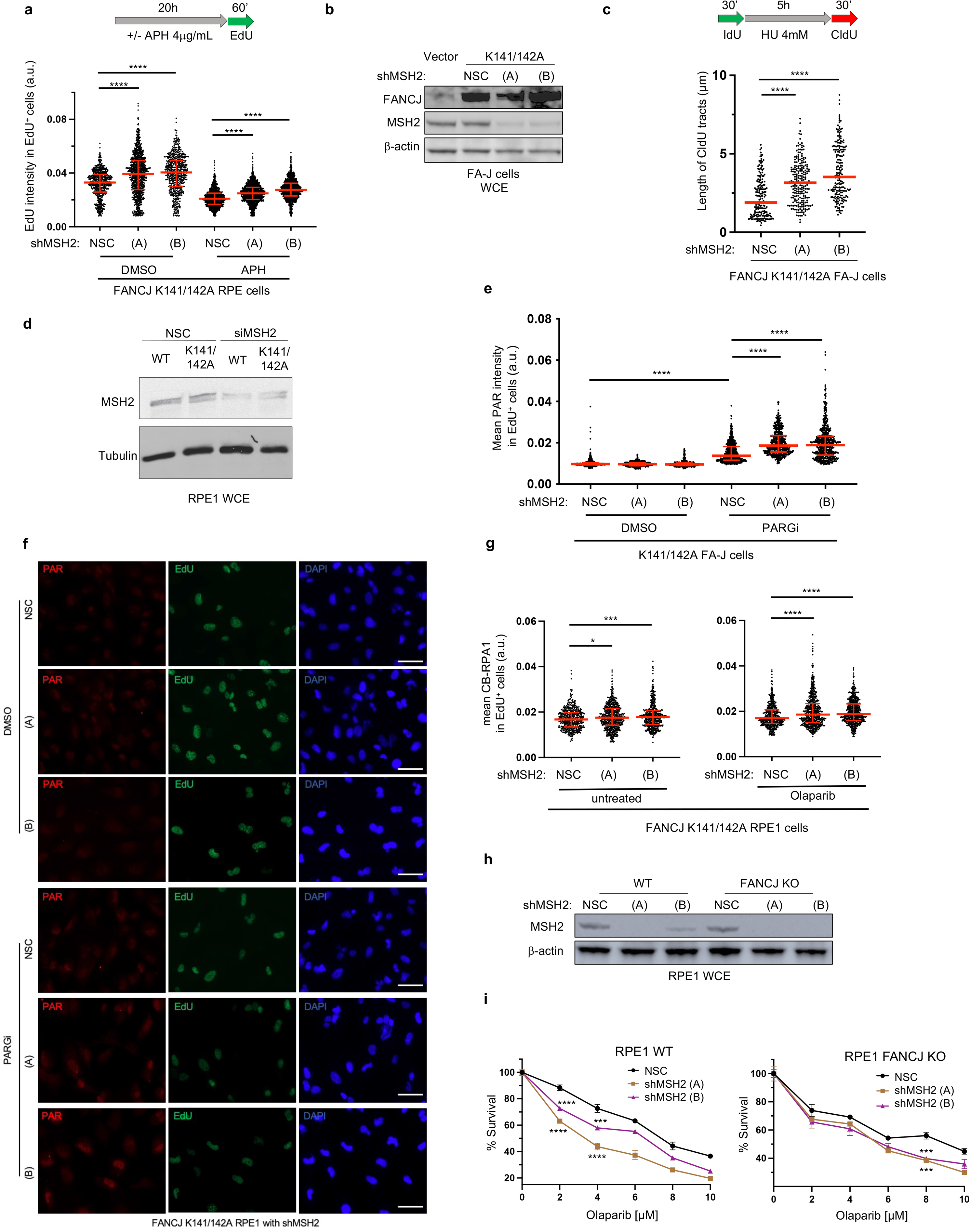
The FANCJ-MLH1 interaction restricts MSH2. **a** Quantification of mean EdU intensity from EdU^+^ cells. EdU was incubated for 1hr after 20 hr DMSO or aphidicolin (APH, 4 µg/ml) treatment in the indicated RPE1 cells. EdU^+^ cells were gated according to positive EdU incorporation. Each dot represents one cell. At least 200 cells are quantified from 2 biological independent experiments (n = 2). Red bars represent the median ± interquartile range. **b-c** WB analysis and quantification of the DNA fiber assays for the length of IdU-connected CldU tracts interrupted by HU treatment (4 mM, 5 h) in FA-J complemented cells with MSH2 depletion, Each dot represents 1 fiber; at least 200 fibers are quantified from n = 2. Red bars represent the median ± interquartile range. **d** WB experiment for whole cell lysates showing the indicated proteins RPE1 WT and mutant cells. **e** Quantification of PAR after 30min DMSO or PARGi (10 µM) treatment in the indicated EdU-positive FA-J cells. **f** Representative images related to Fig 4c. Scale bars, 50 µm **g** Quantification of CB-RPA1 for the indicated cells for (left) untreated or (right) following the Olaparib (10µM, 6h), with EdU incubated for the final 40 min. For the quantification in **e** and **g**, EdU^+^ cells were gated according to positive EdU incorporation. Each dot represents one cell. At least 300 cells are quantified from n = 3. Red bars represent the median ± interquartile range. **h, i** WB analysis and cell survival assays for the indicated RPE1 cells under increasing concentrations of olaparib. Data represent the mean percentage ± SD of survival for each dot. Significance was determined by one-way ANOVA followed by Dunnett’s test comparing NSC to shMSH2 cells. All the other statistical analysis above according to Kruskal-Wallis test, followed by Dunn’s test.

**Supplementary Figure 5:**
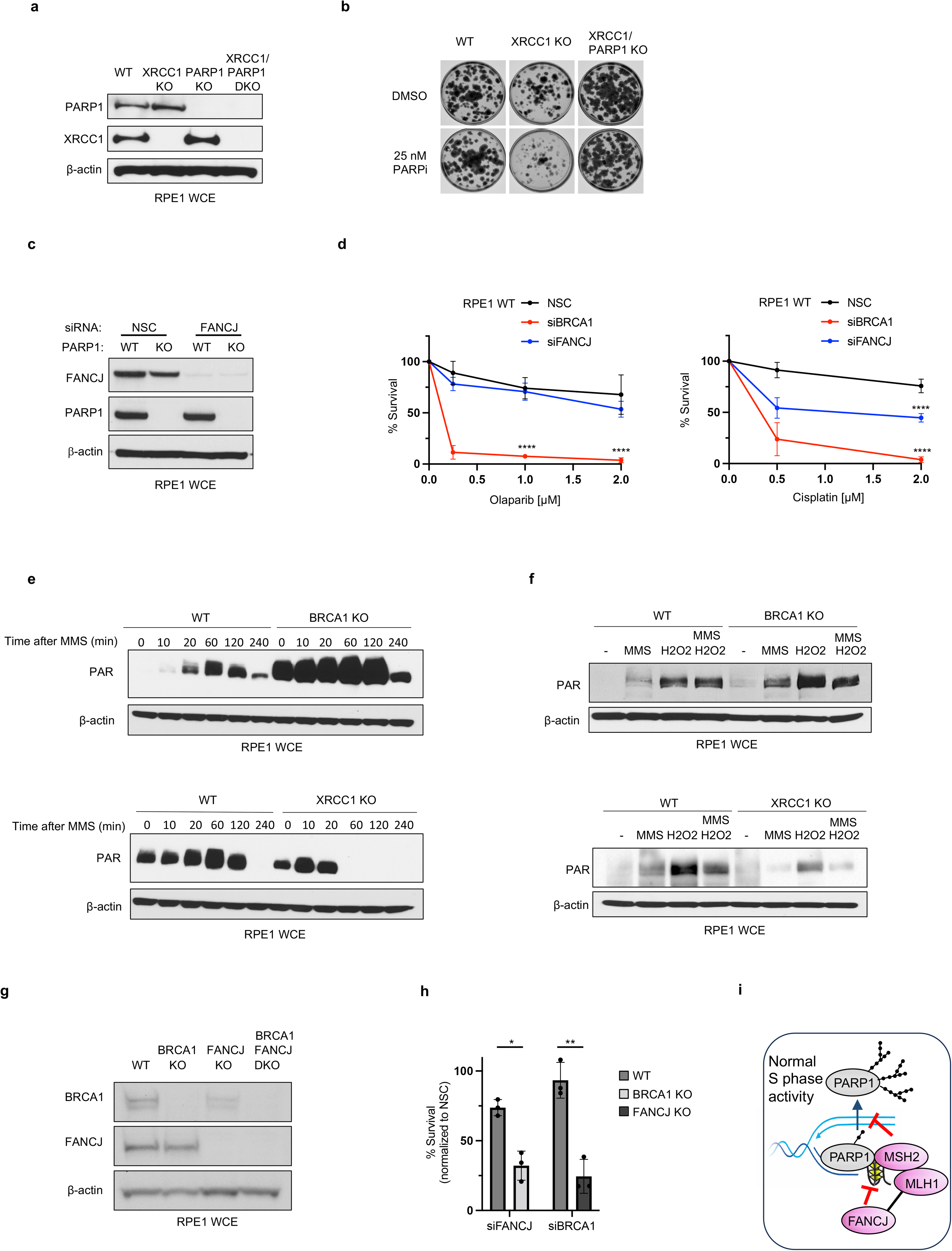
PARP1 deletion rescues chemosensitivity in XRCC1 deficiency without impairing cell fitness. **a** Representative WB for the whole cell lysates showing indicated proteins in untreated WT, XRCC1 KO, PARP1 KO and XRCC1/PARP1 KO RPE1 cells. **b** Clonogenic survival assay of the indicated RPE1 cell lines treated with PARPi. **c** Representative WB for the whole cell lysates showing indicated proteins in untreated WT vs PARP1 KO RPE1 cells with or without FANCJ depletion. **d** Cell survival assays for the indicated cells under increasing concentrations of Olaparib and Cisplatin (CDDP) treatment. Data represent the mean percentage ± SD of survival for each dot. Significance was determined by one-way ANOVA followed by Dunnett’s test comparing NSC to other cells. **e** Representative WB for the whole cell lysates showing PAR in the indicated isogenic cells following MMS (0.1mg/ml) treatment at different time points for 0-240min. **f** Representative WB for the whole cell lysates showing PAR in the indicated isogenic cells following MMS (0.1mg/ml, 60 min), H_2_O_2_ (2mM, 10 min), or combined treatment (MMS 0.1mg/ml 60 min followed by H_2_O_2_ 2mM, 10min). **g** Representative WB for the whole cell lysates for the indicated cell lines and proteins. **h** Quantification of clonogenic survival assays of the indicated RPE1 cell lines and knockdown reagents. Mean survival percentages normalized to non-silencing control (NSC) within each genetic line. n=3 ± SD. Significance determined by one-way ANOVA followed by Dunnett’s test. **i** Model that FANCJ dismantles replisome-associated MSH2-bound G4s through its MLH1 binding and resolves G4 structures to activate and release PARP1 from chromatin.

